# Dopamine receptor DOP-1 engages a sleep pathway to modulate swimming in *C. elegans*

**DOI:** 10.1101/2021.02.17.431715

**Authors:** Ye Xu, Lin Zhang, Yan Liu, Irini Topalidou, Cera Hassinan, Michael Ailion, Zhenqiang Zhao, Tan Wang, Zhibin Chen, Jihong Bai

**Author notes:** Jihong Bai, Ph.D. (Lead Contact and Co-corresponding author) Fred Hutchinson Cancer Research Center 1100 Fairview Ave N. Seattle, WA 98109, USA. Zhibin Chen, M.D., Ph.D., Tan Wang, M.D., Ph.D. (Co-corresponding author) Department of Neurology First Affiliated Hospital of Hainan Medical University Haikou, P. R. China, 570102.

## Abstract

Animals require robust yet flexible programs to support locomotion. While it is clear that a variety of processes must be engaged to ensure rhythmic actions, the exact mechanisms remain largely unknown. Here we report a novel pathway that connects the D1-like dopamine receptor DOP-1 with a sleep mechanism to modulate swimming in *C. elegans*. We show that DOP-1 plays a negative role in sustaining swimming behavior. By contrast, a pathway through the D2-like dopamine receptor DOP-3 negatively regulates the initiation of swimming, but its impact fades quickly over a few minutes. We find that DOP-1 and the GPCR kinase GRK-2 function in the sleep interneuron RIS, where DOP-1 modulates the secretion of a sleep neuropeptide FLP-11. Our genetic studies further show that DOP-1 and FLP-11 act in the same pathway to modulate swimming. Together, these results delineate a functional connection between a dopamine receptor and a sleep program to regulate swimming in *C. elegans*. The temporal transition between DOP-3 and DOP-1 pathways highlights the dynamic nature of neuromodulation for rhythmic movements that persist over time.

**HIGHLIGHTS:**

1. The D1-like dopamine receptor DOP-1 regulates swimming at 10 minutes
2. An integrated function of DOP-1 and DOP-3 is required for the continuity of swimming
3. DOP-1 and GRK-2 act in the sleep neuron RIS
4. FLP-11, a neuropeptide that promotes sleep, negatively regulates swimming

**IN BRIEF:** Xu et al. investigated genetic programs that modulate swimming behavior in the nematode *C. elegans*. They identified a functional link that couples a D1-like dopamine receptor to a sleep program that modulates the sustained phase rather than the initial phase of swimming.

## INTRODUCTION

Animals use various locomotion strategies to benefit their survival. Even simple forms of locomotion like crawling, walking, or swimming require the nervous system to engage muscles in a timely and precise manner. Both invertebrate and vertebrate animals have developed neural circuits to produce stable motor outputs with rhythmic movements (Getting, 1989; Grillner, 2006; Marder and Bucher, 2001; Marder and Calabrese, 1996; Pearson, 2000; Ruder and Arber, 2019). To readily respond to challenges, these neural circuits must be robust yet flexible.

Decades of research show that sustained movements start with activities of central pattern generators (CPGs), components of the nervous system that are capable of producing oscillatory activity without receiving extrinsic information (Delcomyn, 1980; Grillner, 2006; Kiehn, 2006; Marder and Bucher, 2001; Marder et al., 2005). However, to smoothly perform behavioral tasks, a complex network of sensory inputs and feedback mechanisms need to be engaged to shape the activity of CPGs for motor coordination and rhythm modulation (Marder et al., 2015; Marder and Thirumalai, 2002; Miles and Sillar, 2011). The nematode *C. elegans* utilizes undulatory locomotion to crawl and swim (Cohen and Sanders, 2014; Pierce-Shimomura et al., 2008; Zhen and Samuel, 2015). Multiple CPG oscillators, made up of motor neurons in the *C. elegans* ventral nerve cord, generate intrinsic network oscillations for body movements (Fouad et al., 2018; Gao et al., 2018; Wen et al., 2018). While it is clear that signals from upstream interneurons and the electrical coupling within the ventral nerve cord are necessary for coordinating pattern generators to produce robust yet flexible oscillations (Kawano et al., 2011; Wen et al., 2012; Xu et al., 2018), the precise modulatory mechanisms that establish dynamic rhythmicity remain largely unknown.

Dopamine is a neurotransmitter of diverse functions including rhythm modulation in both vertebrate and invertebrate animals (Cermak et al., 2020; Harris-Warrick et al., 1998; Puhl and Mesce, 2008; Sharples et al., 2014; Svensson et al., 2003; Vidal-Gadea et al., 2011). In *C. elegans*, dopamine regulates behaviors such as habituation of mechanical stimuli (Ardiel et al., 2016; Rose and Rankin, 2001; Sanyal et al., 2004), slowing of locomotion in the presence of food (Sawin et al., 2000), the transition from swimming to crawling (Vidal-Gadea et al., 2011), spatial pattern selection (Han et al., 2017), as well as learning and decision making (Calhoun et al., 2015; Rengarajan et al., 2019; Wang et al., 2014). The impact of dopamine on behaviors in *C. elegans* is determined by several factors. First, dopamine receptors are widely expressed in the nervous system (Bentley et al., 2016; Chase et al., 2004; Sugiura et al., 2005; Suo et al., 2002, 2003; Tsalik et al., 2003), resulting in many interaction nodes between dopamine and motor circuits. Second, different dopamine receptors in *C. elegans*, like those in vertebrates, have distinct functions. For instance, in worms, the D1-like dopamine receptor DOP-1 and the D2-like dopamine receptor DOP-3 have opposite effects on locomotion (Chase et al., 2004) and acetylcholine release (Allen et al., 2011), likely through cell-autonomous mechanisms in motor neurons. Third, the strength of dopamine signals through receptor-mediated responses is tuned by G proteins and their regulatory mechanisms (Jiang et al., 2001; Wang et al., 1995). In the case of basal slowing of locomotion on food, DOP-3 activates the Gαo pathway, while DOP-1 acts through the Gαq signaling pathway (Chase et al., 2004). Recently, Topalidou et al. identified a novel mechanism at the interface between locomotion and dopamine signaling in which the G protein coupled receptor kinase-2 (GRK-2) negatively modulates a DOP-3 pathway in premotor interneurons (Topalidou et al., 2017b). GRK-2 leads to inactivation of GOA-1 and activation of NALCN/NCA channels, which subsequently support rhythmic outputs of body bends during crawling and swimming.

Neuropeptides are also key regulators of locomotory activity. Their action to inhibit or activate locomotion is often linked to developmental clocks (Erion et al., 2016; Krupp et al., 2013; Selcho et al., 2017; Taghert and Nitabach, 2012; Xu et al., 2008). During development, *C. elegans* undergoes four larval molts with a cycle of approximately 8-10 hours (Moss, 2007). During each larval molt, the worms enter into a prolonged period (∼2 hours) of behavioral quiescence with inactivation of locomotion and feeding behaviors, with properties similar to the sleep-like state in mammals and *Drosophila melanogaster* (Hendricks et al., 2000; Huber et al., 2004; Nelson and Raizen, 2013; Raizen et al., 2008; Shaw et al., 2000). Neuropeptide pathways have been found to regulate this developmentally-timed behavior quiescence in *C. elegans* (Choi et al., 2013; Turek et al., 2016; Van der Auwera et al., 2020). Among them, a FMRFamide-related neuropeptide FLP-11, stored in the sleep interneuron RIS, strongly inhibits locomotion at the onset of sleep (Turek et al., 2016).

In this study, we report a functional link between the dopamine receptor DOP-1 and the sleep peptide FLP-11 for sustaining swimming behavior. We find that DOP-1 and DOP-3 negatively regulate *C. elegans* swimming through two different circuits. While both pathways are responsible for the defective swimming pattern of *grk-2* mutants, the impact of DOP-1 and DOP-3 on swimming is restricted to distinct temporal periods. Together, these results demonstrate a functional integration of a dopamine receptor and a sleep quiescence program in living animals, revealing the dynamic nature of modulation underlying the seemingly constant appearance of repetitive movements.

## RESULTS

### A requirement for different signals during early and late stages of swimming

We quantified the swimming frequency by tracing changes of mid-point body angle over time. As previously reported (Topalidou et al., 2017b), *grk-2* mutants exhibited severe defects in swimming behavior (Figure 1), supporting a positive role of GRK-2 in modulating swimming. The body bend frequency of *grk-2* mutants was decreased to less than 0.5 Hz from approximately 2 Hz in wild type animals. The reduction in swimming frequency in *grk-2* mutants was largely suppressed by a *dop-3(vs106) null* mutation (Figure 1A, C). These results are consistent with previous findings showing that DOP-3 is a negative regulator of swimming (Topalidou et al., 2017b). However, at 10 min, the body bend frequency of the *grk-2; dop-3* double mutant was significantly decreased to a level (∼0.4 Hz) that is indistinguishable from the frequency of *grk-2* single mutant swimming (Figure 1B, C). Indeed, more than 60% of the *grk-2; dop-3* mutants completely stopped swimming by 10 min (Figure 1C right panel). These results showed that the *dop-3* mutation suppresses the swimming defect of *grk-2* mutants at 1-minute but not at 10-minutes. The reduction of swimming frequency was unlikely due to osmolarity changes upon transferring worms from NGM plates to M9 buffer, as *grk-2; dop-3* mutants showed similar decreases in swimming frequency in NGM salt solution (See also Figure S1A).

**Figure 1.**
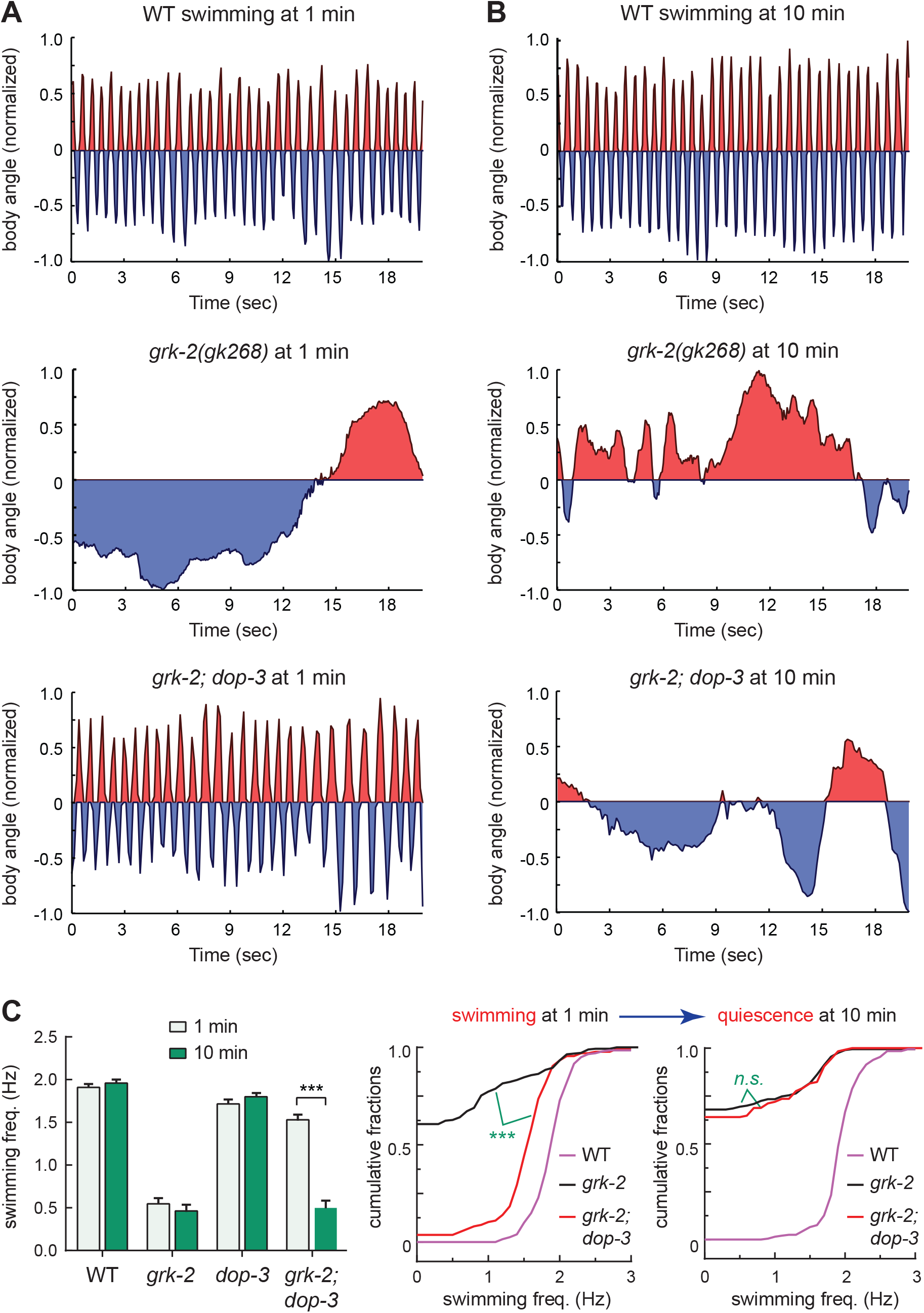
The *grk-2; dop-3* double mutant worms failed to sustain swimming. Changes of mid-point body angles were plotted over a time window of 20 seconds during swimming. For clarity, positive angles are indicated in red and negative angles are indicated in blue. (A) Worms were placed in M9 buffer for 1 minute. Representative swimming traces are shown for wild type (N2; *top panel*), *grk-2(gk268)* single mutant *(middle panel),* and *grk-2(gk268); dop-3(vs106)* double mutant worms *(lower panel)*. (B) Worms were placed in M9 buffer for 10 minutes. Representative swimming traces show that *grk-2; dop-3* double mutants stopped swimming at the 10 min time point. (C) Quantification of data. *Left:* Body bend frequency was quantified and plotted as mean ± SEM for N2, *grk-2(gk268)*, *dop-3(vs106)*, and *grk-2(gk268); dop-3(vs106)* mutants. The *grk-2(gk268); dop-3(vs106)* double mutants exhibited a swimming defect at the 10 min time point. \*\*\**p < 0.001*, (unpaired Student’s t-test). *Middle and right panels:* Cumulative fraction plots of swimming frequency distribution at 1 min (*middle*) and 10 min (*right*) time points. Swimming of *grk-2(gk268); dop-3(vs106)* double mutants exhibited a shift from high frequency at 1 min to low frequency at 10 min. The frequency distribution curves of *grk-2* single and *grk-2; dop-3* double mutants were significantly different at 1 min, but they became identical at 10 min (Kolmogorov-Smirnov test). \*\*\**p < 0.001*, *n.s.,* not significant. Statistical analysis and graphing were carried out using Prism 8 (GraphPad).

To determine whether the *grk-2; dop-3* mutants had lost their physical ability to swim at 10-min, we poked these worms using a platinum wire. We found that more than 90% of the worms bended their body immediately after poking (See also Figure S1B). However, when examined at 30 seconds after poking, about 70% of the worms had stopped swimming (See also Figure S1B). These data indicate that the *grk-2; dop-3* worms at 10-min were in a quiescent rather than a paralyzed state, as they could initiate but were unable to sustain the body bend activity. Interestingly, after entering the quiescent state, the *grk-2; dop-3* mutants stopped pharyngeal pumping (See also SI video 1), similar to worms undergoing lethargus on food (Raizen et al., 2008; Van Buskirk and Sternberg, 2007), but different from those experiencing exercise-induced quiescence (Schuch et al., 2020).

Together, these results show that the *grk-2; dop-3* mutants enter a quiescent state at 10-min, indicating that the impact of the D2-like dopamine receptor DOP-3 on swimming fades quickly, and additional mechanisms are necessary for sustained swimming. In contrast to *dop-3(vs106)*, a *cat-2(e1112)* mutation that disrupts tyrosine hydroxylase for dopamine synthesis suppressed swimming defects in *grk-2* mutants at both 1 min and 10 min (See also Figure S1C), suggesting that swimming at 10-minutes is regulated by dopamine through receptors other than DOP-3.

### DOP-1 and DOP-3 act in distinct phases of swimming

*C. elegans* locomotion is modulated by dopamine through the antagonistic action of the dopamine receptors DOP-1 and DOP-3 (Chase et al., 2004). To determine whether DOP-1 plays a role in delayed swimming quiescence, we introduced the *dop-1(vs100)* mutation into *grk-2; dop-3* mutant worms, which can initiate but fail to sustain swimming. We found that *grk-2; dop-1 dop-3* triple mutants remained active swimmers after 10 minutes (Figure 2A). Body bend frequency of *grk-2; dop-1 dop-3* triple mutants was identical at 1 min and 10 min (Figure 2B), and the cumulative distribution of the swimming frequency was shifted to significantly higher values in the *grk-2; dop-1 dop-3* triple mutants compared to the *grk-2; dop-3* double mutants (Figure 2C). As a control, we investigated the role of DOP-4, the other D1-like dopamine receptor in *C. elegans*. We found that the *dop-4(tm1392)* mutant did not restore swimming of *grk-2; dop-3* mutants at 10 min (See also Figure S2). To further confirm the role of *dop-1*, we examined the swimming of *grk-2; dop-1* double mutants. We found that these mutant worms were more active in swimming at 10-min than they were at 1-min, indicating a late role of DOP-1 in inhibiting swimming (Figure 3A). Together, these data show that both DOP-1 and DOP-3 are negative regulators of swimming, but their roles are distributed to distinct phases of swimming: DOP-3 modulates early swimming while DOP-1 regulates the late phase of swimming.

**Figure 2.**
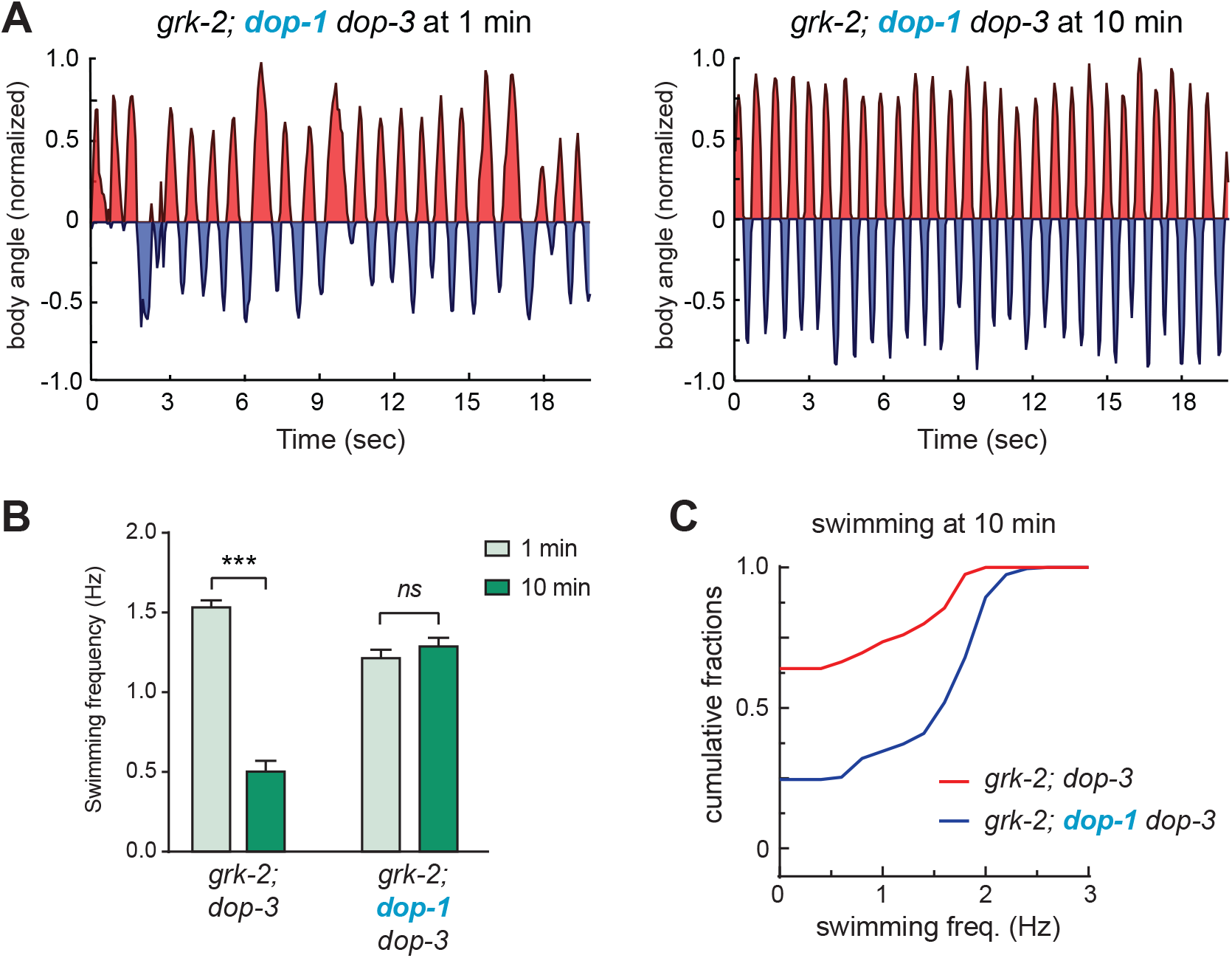
DOP-1 and DOP-3 support distinct phases of swimming. Representative swimming traces are shown for *grk-2(gk268); dop-1(vs100) dop-3(vs106)* triple mutants (A). *Left panel*, swimming at 1 min. *Right panel*, swimming at 10 min. (B) Body bend frequency was plotted as mean ± SEM. *grk-2; dop-1 dop-3* triple mutants showed identical swimming frequencies at 1 min and 10 min. \*\*\**p < 0.001*, (unpaired Student’s t-test). *n.s.,* not significant. (C) Cumulative fraction plots of swimming frequency distribution show a significant difference between *grk-2; dop-3* double and *grk-2; dop-1 dop-3* triple mutants at 10 min (Kolmogorov-Smirnov test).

**Figure 3.**
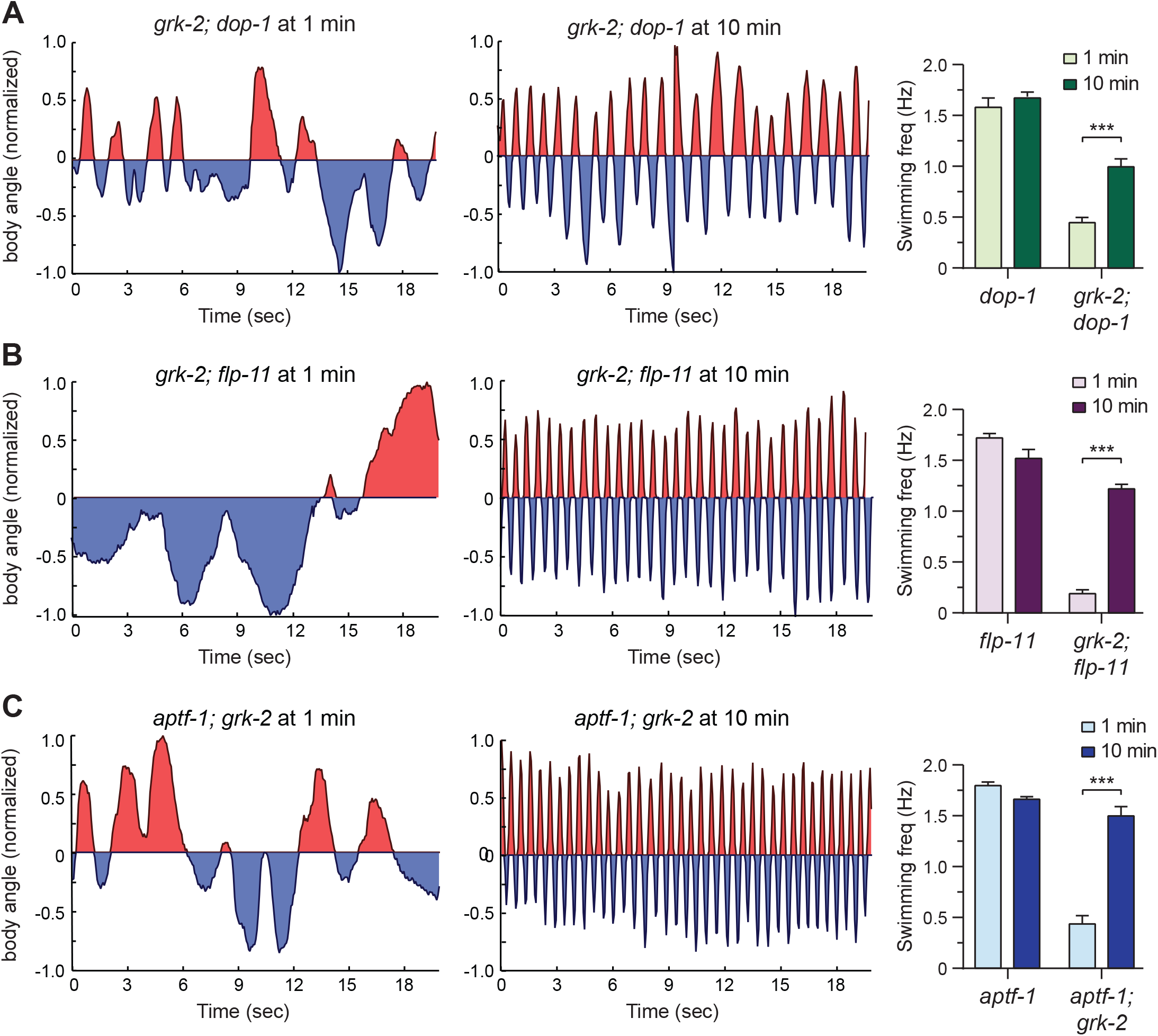
DOP-1, neuropeptide FLP-11 and APTF-1 modulate swimming at 10-minutes. (A) Representative traces show the swimming behavior of *grk-2(gk268); dop-1(vs100)* double mutants at 1 min (*left*) and 10 min (*middle*). Body bend frequency was quantified and plotted in the *right* panel. (B) Representative traces show the swimming behavior of *grk-2(gk268); flp-11(tm2706)* double mutants at 1 min (*left*) and 10 min (*middle*). Body bend frequency was quantified and plotted in the *right* panel. (C) Representative traces show the swimming behavior of *aptf-1(gk794); grk-2(gk268)* double mutants at 1 min (*left*) and 10 min (*middle*). Body bend frequency was quantified and plotted as mean ± SEM (*right*). \*\*\**p < 0.001*, (unpaired Student’s t-test). Statistical analysis and graphing were carried out using Prism 8 (GraphPad).

### The sleep neuropeptide FLP-11 regulates swimming at a late-phase

To gain insight into the mechanisms behind the late-phase quiescence, we looked into pathways that are capable of inducing locomotion quiescence. In particular, we investigated several genes encoding neuropeptides that reduce locomotion activity during sleep-like states. We created double mutants between *grk-2* and individual mutations that disrupt *flp-11, flp-18, flp-21,* or *pdf-1* (Bhardwaj et al., 2018; Choi et al., 2013; Turek et al., 2016). We found that *grk-2; flp-11* double mutants exhibited strong recovery of swimming at 10 min (Figure 3B). By contrast, disruption of *flp-18, pdf-1,* or *flp-21* did not restore swimming in *grk-2* mutants (See also Figure S3). Finally, removal of the AP2 transcription factor APTF-1, which controls the expression of FLP-11 (Turek et al., 2016; Turek et al., 2013), induced a significant recovery of swimming in *grk-2* mutants at 10 minutes, which is similar to the swimming recovery in *grk-2; flp-11* double mutants (Figure 3C). Together, these findings indicate that the FLP-11 pathway plays a role in the late stage of swimming.

### DOP-3 and FLP-11 pathways act independently

Because *grk-2; flp-11* and *grk-2; dop-3* worms swim at different time points, we hypothesized that the FLP-11 and DOP-3 pathways function in independently continuous swimming. In support of this idea, simultaneous removal of both *flp-11* and *dop-3* from *grk-2* mutants led to recovered swimming at both 1 and 10 minutes with identical frequencies (Figure 4A; 1 min: 1.02 ± 0.04 Hz; and 10 min: 0.94 ± 0.04 Hz; *n.s.*), similar to *grk-2; dop-1 dop-3* mutants. The additive impact of DOP-3 and FLP-11 on the sustainability of swimming supports the idea that these genetic pathways act in distinct phases of swimming. Similar to the *dop-1* mutation (Figure 2B), deletion of *flp-11* from *grk-2; dop-3* double mutants also caused a minor reduction in the rate of body bends at 1-min (Figure 4A), suggesting that FLP-11 and DOP-1 have an additional role in supporting early swimming, the opposite of their role in regulating swimming at 10 minutes.

**Figure 4.**
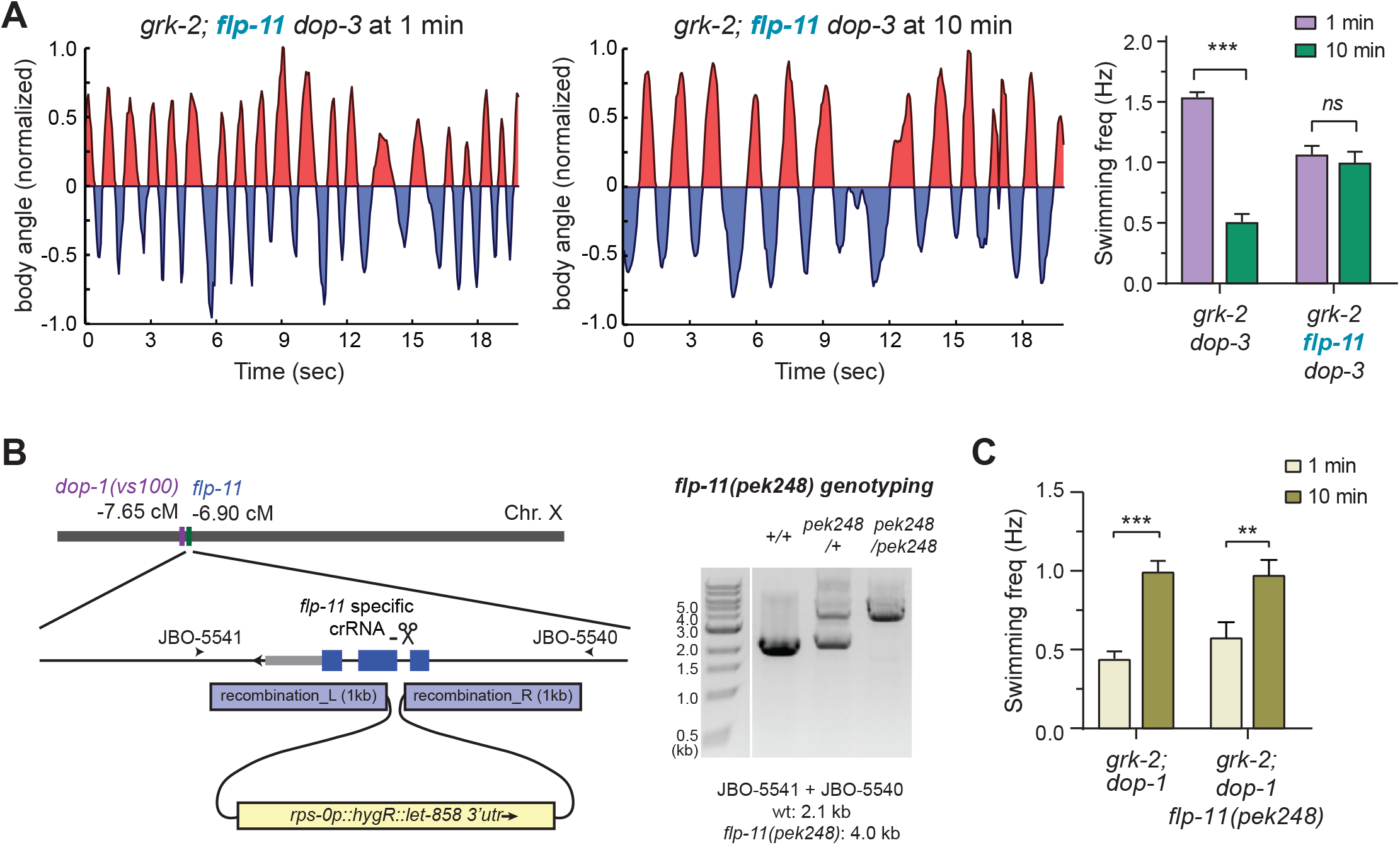
*dop-1,* but not *dop-3*, functions in the same genetic pathway with *flp-11*. (A) Representative traces show the swimming behavior of *grk-2(gk268); flp-11(tm2706) dop-3(vs106)* triple mutants at 1 min (*left*) and 10 min (*middle*). Body bend frequency was quantified and shown as mean ± SEM *(right)*. \*\*\**p* < 0.001, (unpaired Student’s t-test). *n.s.,* not significant. (B) The *left* panel shows a schematic diagram for the CRISPR-Cas9 and homologous recombination strategy used to disrupt the *flp-11* gene in *grk-2; dop-1* mutants. The genotype of the *flp-11*(*pek248*) mutant allele was confirmed by PCR amplification using oligos JBO-5540 and JBO-5541. Genotyping results are shown in the *right* panel. (C) Body bend frequency was analyzed for *grk-2(gk268); dop-1(vs100)* double mutants and *grk-2(gk268); dop-1(vs100) flp-11(pek248)* triple mutants, and was plotted as mean ± SEM. \*\*\**p < 0.001*, \*\**p < 0.01*, (unpaired Student’s t-test). Statistical analysis and graphing were carried out using Prism 8 (GraphPad).

### DOP-1 and FLP-11 act in the same genetic pathway

Next, we examined the genetic relationship between *dop-1* and *flp-11*. Because *flp-11* and *dop-1* are located close to each other (∼ 0.7cM) on chromosome X, we employed a CRISPR-Cas9 and homologous recombination strategy to directly disrupt the *flp-11* gene in *grk-2; dop-1* mutants (Figure 4B). We obtained a *flp-11*(*pek248*) mutant allele that carries a deletion within the 1^st^ intron and the 2^nd^ exon of *flp-11* and a large insertion (1.9kb) of a hygromycin-resistance (HygR) cassette in the 2^nd^ exon (Figure 4B). Because these changes disrupt the gene structure of *flp-11*, we expect that the *pek248* mutation is a null allele. Introduction of the *flp-11*(*pek248*) mutation into *grk-2; dop-1* worms did not further enhance the phenotype of *grk-2; dop-1* mutants (Figure 4C), indicating that *flp-11* and *dop-1* function in the same genetic pathway.

### Premotor interneurons and the sleep neuron RIS regulate different phases of swimming

GRK-2 is expressed in many neurons including acetylcholine head neurons and body motor neurons (Topalidou et al., 2017b). Since expression of GRK-2 in premotor interneurons significantly restores locomotion activity during crawling (Topalidou et al., 2017b), we asked whether swimming is also controlled by GRK-2 in premotor interneurons. We introduced a single-copy transgene of *nmr-1p::grk-2* into *grk-2* mutants. These worms were fully active in early swimming (1 min) with a frequency of 1.7 ± 0.07 Hz (Figure 5), indicating a positive role of GRK-2 in premotor interneurons to initiate swimming. However, the swimming frequency of these worms was significantly reduced to 1.1 ± 0.11 Hz by 10 min, demonstrating that GRK-2 in premotor interneurons could not fully sustain swimming. Therefore, additional neurons must be involved to ensure the continuity of swimming.

**Figure 5.**
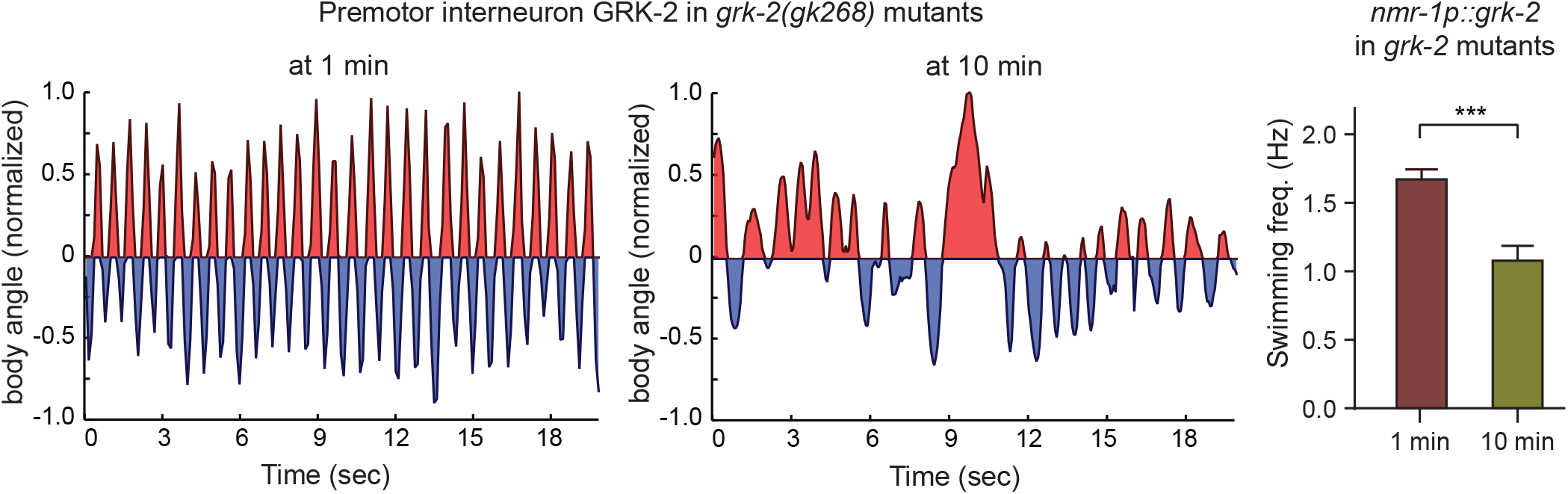
GRK-2 acts in premotor interneurons to modulate early phase of swimming. Representative traces show the swimming behavior of transgenic *grk-2(gk268)* worms carrying a single-copy *nmr-1p::grk-2* transgene at 1 min (*left*) and 10 min (*middle*), respectively. Body bend frequency was plotted as mean ± SEM (*right*). \*\*\**p < 0.001*, (unpaired Student’s t-test). Statistical analysis and graphing were carried out using Prism 8 (GraphPad).

FLP-11 is secreted from the sleep neuron RIS to inhibit locomotion during development (Steuer Costa et al., 2019; Turek et al., 2016). Consistent with previous findings, we found that the *flp-11* promoter (*flp-11p*) drove specific expression in the RIS neuron (See also Figure S4A). The tight pattern of *flp-11* expression prompted us to determine whether the RIS sleep neuron is the functional site for GRK-2 and DOP-1.

First, we asked whether *grk-2* is expressed in the RIS neuron. We generated transgenic worms carrying a nuclear green fluorescence reporter 2xNLS::mNeonGreen in RIS (*flp-11p::2xnls::mNeongreen*) and a cytoplasmic red fluorescence reporter TagRFP under the control of *grk-2p* (*grk-2p::tagrfp*). Using confocal fluorescence microscopy, we found that the RIS neuron had both nuclear green and cytoplasmic red fluorescence (Figure 6A), indicating that GRK-2 is expressed in RIS, supporting previous observations through transcriptome analysis (Konietzka et al., 2020).

**Figure 6.**
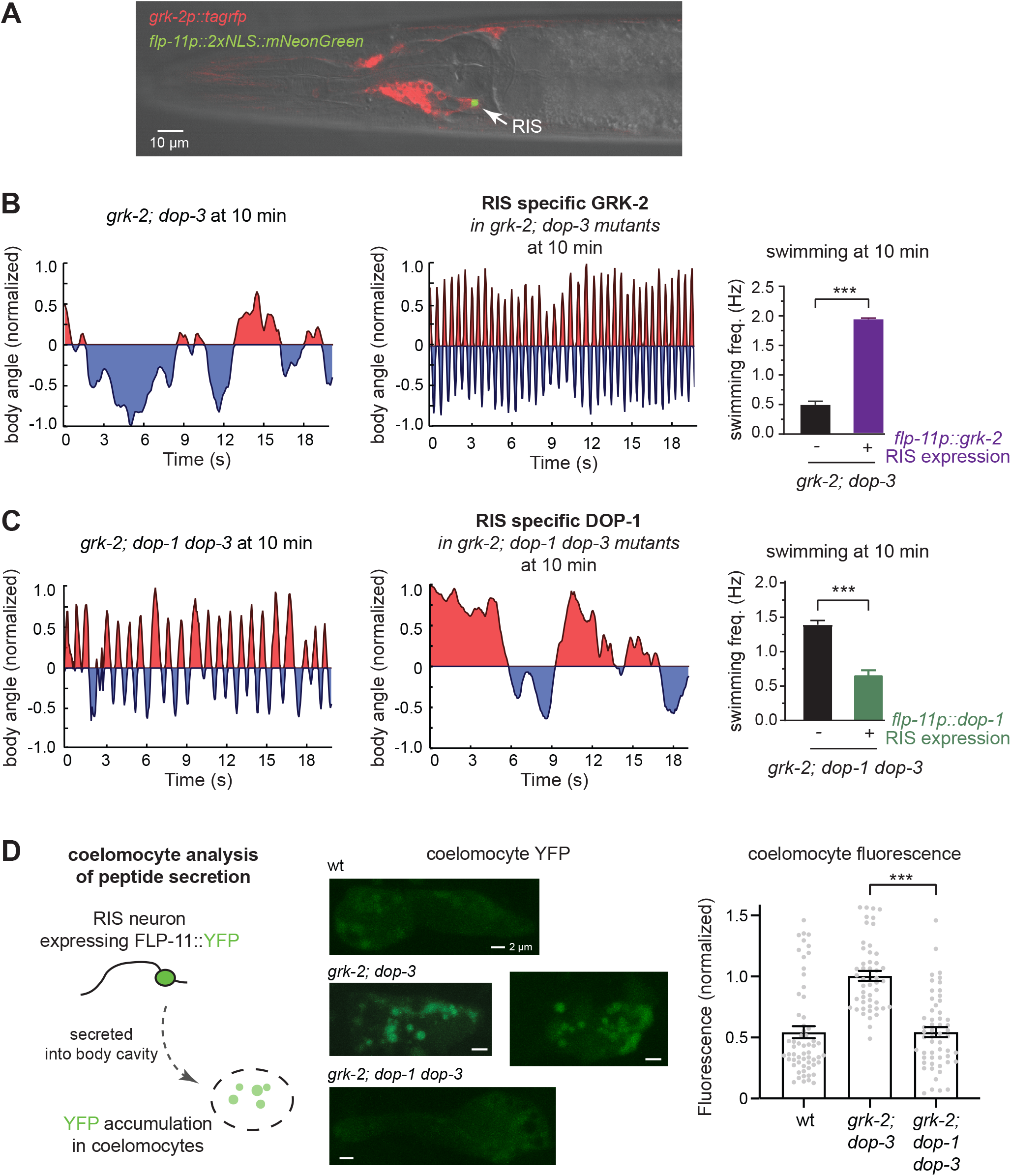
The sleep interneuron RIS is the functional site for GRK-2 and DOP-1 to regulate late-stage swimming. (A) An image showing GRK-2 expression in the RIS neuron. Transgenic worms carrying *grk-2p::tagrfp* and *flp-11p::2xnls::mNeonGreen* were examined using an Olympus FV1000 scanning confocal microscope. The green fluorescent protein mNeonGreen was excited by a 488 nm laser, and the red fluorescent protein TagRFP was excited using a 559 nm laser. The 559 nm laser was also used to collect DIC (differential interference contrast) images. The fluorescent image was superimposed on the DIC image for clarity. (B) Representative swimming traces for *grk-2(gk268); dop-3(vs106)* double mutants, in the presence (*left*) or absence (*middle*) of a single-copy transgene *flp-11p::grk-2*. Body bend frequency at 10 min was quantified and shown as mean ± SEM in the *right* panel. (C) Representative swimming traces for *grk-2(gk268); dop-1(vs100) dop-3(vs106)* triple mutants, in the presence (*left*) or absence (*middle*) of a *flp-11p::dop-1* transgene. Body bend frequency at 10-min was quantified and shown as mean ± SEM (*right*). (D) The *left* panel shows a schematic diagram of the coelomocyte assay used to quantify FLP-11 secretion from RIS. Worms carrying an integrated transgene (*pekIs116 [flp-11p::flp-11::yfp]*) were examined using the Olympus FV1000 scanning confocal microscope. The yellow fluorescent protein mVenus was excited using the 488 nm laser. Representative images of anterior coelomocytes in wild type (wt), *grk-2(gk268); dop-3(vs106)* double, and *grk-2(gk268); dop-1(vs100) dop-3(vs106)* triple mutant worms are shown in the *middle*. Fluorescence intensity of YFP puncta in coelomocytes was quantified and is shown in the *right* panel. \*\*\**p < 0.001*, (one-way ANOVA with Dunnett’s multiple comparisons test).

Next, to determine whether GRK-2 acts in RIS to sustain swimming, we expressed a *flp-11p::grk-2* transgene in *grk-2; dop-3* worms that are able to initiate swimming but fail to sustain it at late stages (Figure 1), and examined the ability of these transgenic worms to swim at the 10-minute time point. In contrast to *grk-2; dop-3* worms that fail to sustain swimming, expression of *flp-11p::grk-2* fully restored the body-bend frequency of *grk-2; dop-3* worms at 10 minutes (Figure 6B). These data show that GRK-2 in the RIS sleep neuron is sufficient to support late-stage swimming. As a control, GRK-2 expression in the premotor interneurons by *nmr-1p* did not show rescue activity at 10-min (See also Figure S4B,C), confirming that the role of premotor neuron GRK-2 is mainly limited to the early-phase swimming. Together, these results indicate that RIS and premotor interneurons are two sites where GRK-2 functions to support swimming at distinct temporal phases.

Finally, we examined the role of DOP-1 in RIS using cell-specific rescue experiments. DOP-1, like GRK-2, is expressed in many neurons including RIS (Konietzka et al., 2020; Tsalik et al., 2003). We found that expression of a *flp-11p::dop-1* transgene in *grk-2; dop-1 dop-3* mutants inhibited late-stage swimming (Figure 6C), indicating that DOP-1 in RIS is sufficient to negatively modulate late-stage swimming. In addition, the *flp-11p::dop-1* transgene slightly increased the swimming frequency at 1 min (See also Figure S4D), suggesting that DOP-1 may have a minor role in RIS to promote early swimming. In summary, these cell-specific rescue experiments show that GRK-2 and DOP-1 act in RIS to modulate late-phase swimming.

### DOP-1 regulates FLP-11 secretion from the RIS neuron

To understand the functional connection between DOP-1 and FLP-11 in RIS, we next quantified the levels of FLP-11 secretion using an established coelomocyte uptake assay (Ch’ng et al., 2008; Fares and Greenwald, 2001; Sieburth et al., 2007). FLP-11::YFP was expressed in RIS using the *flp-11* promoter. We quantified YFP fluorescence intensity in coelomocytes as a readout of secreted FLP-11::YFP that had been taken up by the scavenger coelomocytes. Our results showed a low level of yellow fluorescence in coelomocytes in wild type worms (Figure 6D), supporting the idea that there is little FLP-11 secretion in awake wild type worms (Turek et al., 2016). By contrast, significantly increased levels of YFP signal were found in *grk-2; dop-3* coelomocytes (Figure 6D), suggesting that FLP-11 secretion was elevated in these double mutants. Finally, we found that, in *grk-2; dop-1 dop-3* triple mutants, coelomocyte YFP fluorescence was returned to a low level (Figure 6D), similar to that in wild type animals. These data show that the secretion of the sleep peptide FLP-11 from the RIS neuron was promoted by the dopamine receptor DOP-1. Because RIS is a sleep neuron and its activity is regulated, we speculated that uncoupling FLP-11 secretion from RIS activity could cause irregularity of swimming activity. Consistent with this idea, we found that ectopic expression of FLP-11 in body wall muscles of wild type worms slightly but significantly reduced swimming frequency at both 1-min and 10-min (See also Figure S4E), suggesting that miscontrolled FLP-11 secretion inhibits swimming.

### Neuropeptide receptors couple FLP-11 to swimming quiescence

As a neuropeptide, FLP-11 requires G-protein-coupled receptors to carry out its function. Previous studies have identified FRPR-3, NPR-4, and NPR-22 as candidate receptors for FLP-11 (Chew et al., 2018; Cohen et al., 2009; Frooninckx et al., 2012; Mertens et al., 2006; Mertens et al., 2004). To determine which neuropeptide receptor acts in the FLP-11 pathway to regulate late-phase swimming quiescence, we examined *grk-2; frpr-3* double mutant worms, and found that removal of FRPR-3 led to a partial but significant increase in swimming frequency at 10 minutes (Figure 7A). Both *npr-22* and *npr-4* mutations, when introduced into *grk-2; frpr-3* mutants, each further enhanced the restoration of swimming (Figure 7A). Finally, the *grk-2; npr-22; frpr-3; npr-4* quadruple mutants showed restored swimming frequency that was indistinguishable from *grk-2; flp-11* double mutants at 10 min (Figure 7A-B). These data show that FLP-11 engages several receptors, whose activation reduces swimming at a late phase.

**Figure 7.**
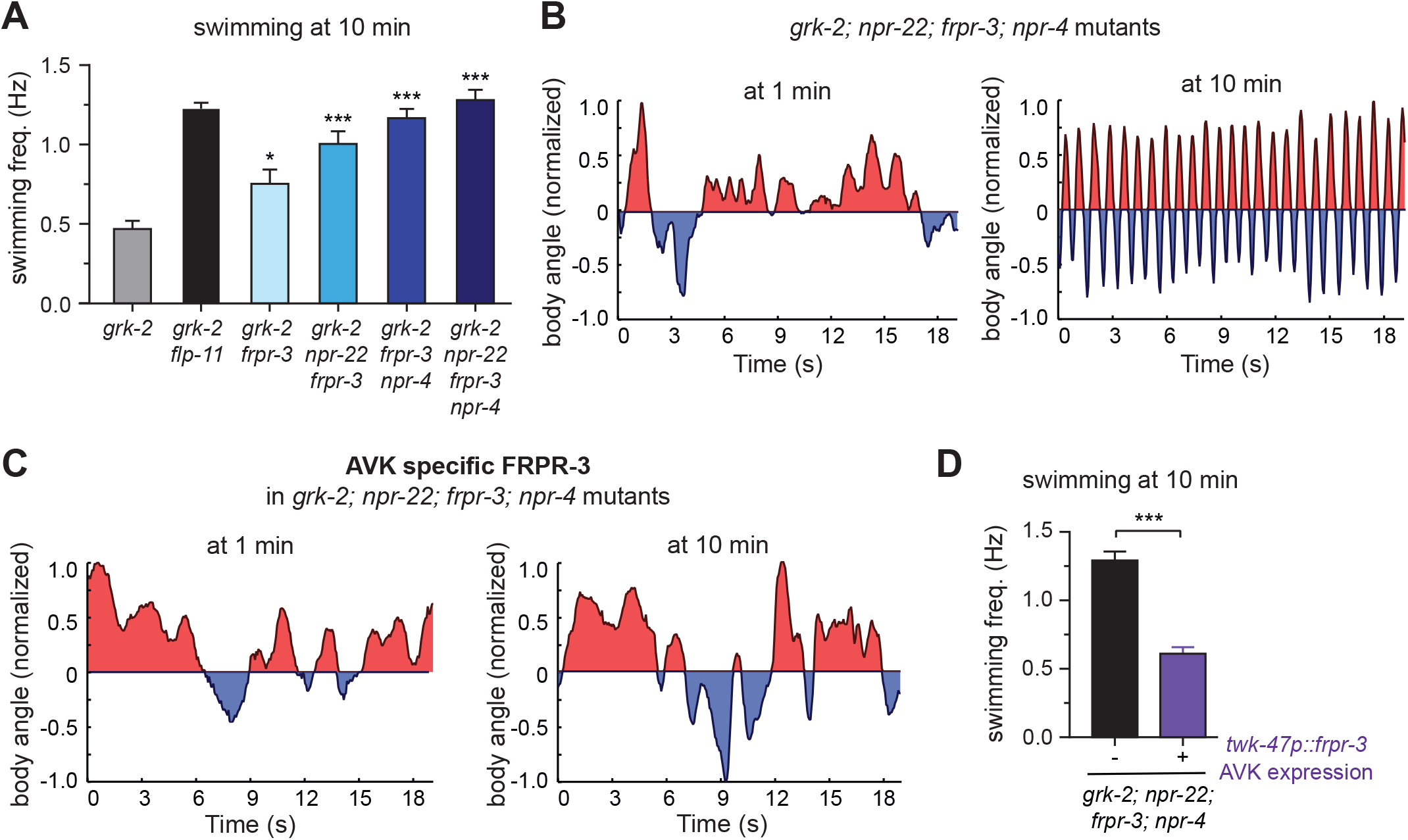
Multiple receptors are responsible for coupling FLP-11 to swimming regulation. (A) Body bend frequency of mutant worms that carry *grk-2(gk268)* and mutations disrupting candidate FLP-11 receptors FRPR-3, NPR-22, and NPR-4 was determined and plotted as mean ± SEM. \**p* < 0.05, \*\*\**p* < 0.001, compared to *grk-2* single mutants (one-way ANOVA with Dunnett’s multiple comparisons test). (B) Representative swimming traces of *grk-2; npr-22; frpr-3; npr-4* mutants at 1 min (*left*) and 10 min (*right*). (C) Representative swimming traces of transgenic *grk-2; npr-22; frpr-3; npr-4* mutant worms that carry *twk-47p::frpr-3* at 1 min (*left*) and 10 min (*right*). (D) Body bend frequency at 10 min; mean ± SEM. \*\*\**p* < 0.001.

Expression of FRPR-3, NPR-4, and NPR-22 is fairly broad in *C. elegans* (Chew et al., 2018; Cohen et al., 2009; Turek et al., 2016). For example, FRPR-3 is expressed in approximately 30 neurons, and NPR-4 is expressed in about 10 neurons. This complex pattern of expression makes it difficult to pinpoint their site-of-action. To get a glimpse into where these neuropeptide receptors function, we investigated the interneuron AVK, which expresses FRPR-3, and receives direct synaptic input from RIS. Previous studies have shown that inhibition of AVK induces slowed swimming, as well as frequent and prolonged stops during crawling (Hums et al., 2016; Oranth et al., 2018). To determine the role of AVK FRPR-3 in swimming, we expressed FRPR-3 using the AVK specific promoter *twk-47p* (Lorenzo et al., 2020) in *grk-2; npr-22; frpr-3; npr-4* mutants. We found that expression of FRPR-3 in AVK significantly reduced swimming frequency at 10 min (Figure 7C-D). These data indicate that expression of FRPR-3 in AVK interneurons is sufficient to induce quiescence during late-stage swimming, suggesting that AVK is one of the neurons that receive FLP-11 signals from RIS to regulate swimming.

## DISCUSSION

We showed that swimming of *C. elegans* is controlled by two distinct dopaminergic pathways. In particular, we found that the D1-like dopamine receptor DOP-1 negatively regulates the frequency of body bends through FLP-11, a neuropeptide required for behavior quiescence during development. We showed that the DOP-1/FLP-11 pathway acts together with the known DOP-3/NALCN regulatory pathway to ensure a continuous pattern of swimming (Pierce-Shimomura et al., 2008; Topalidou et al., 2017b). While both pathways regulate swimming, their action differs. Physiologically, the DOP-3/NALCN pathway functions early during an initial phase of swimming. By contrast, the DOP-1/FLP-11 regulation acts late to inhibit swimming. The DOP-1/FLP-11 and DOP-3/NALCN pathways also differ at the circuit level. The DOP-3/NALCN pathway functions in premotor interneurons (Topalidou et al., 2017b). However, DOP-1 and FLP-11 act in the RIS sleep interneuron. Together, these results show that orchestration of dopaminergic and neuropeptidergic signals sets the foundation for sustained swimming.

### An integrated dopamine signaling network for distinct phases of rhythmic behavior

Swimming is a classic example of a rhythmic behavior that requires neurons to produce locomotor patterns. Unlike reflexes, body bends during swimming occur in continuous sequences in a repeated manner. In *C. elegans*, dopamine plays a key role in modulating several aspects of swimming. For instance, dopamine is necessary and sufficient for worms to switch locomotory patterns from swimming to crawling, and such transition of locomotory gaits requires D1-like dopamine receptors (DOP-1 and DOP-4), but not D2-like receptors (DOP-2 and DOP-3) (Vidal-Gadea et al., 2011). In addition, worms that lack the plasma membrane dopamine transporter DAT-1, when placed in water, swim briefly and then become paralyzed (termed <u>sw</u>im-induced <u>p</u>aralysis; SWIP)(Hardaway et al., 2012; McDonald et al., 2007). While SWIP and the swimming quiescence of *grk-2; dop-3* double mutants appear to be phenotypically similar, the underlying mechanisms are different – SWIP depends solely on DOP-3 (McDonald et al., 2007), which is absent in the *grk-2; dop-3* mutant worms. Instead, the swimming quiescence of *grk-2; dop-3* mutants is mediated by DOP-1 and FLP-11, a neuropeptide secreted from the sleep neuron RIS. Unlike the swim-to-crawl transition, DOP-4 is not required for the swimming quiescence observed in our study. Together, *C. elegans* swimming is modulated by a network of distinct dopaminergic pathways.

While the frequency of movements remains largely constant over a long period of swimming, mechanisms for sustaining the rhythm appear to be dynamic. The impact of the DOP-3 pathway is at the initial phase of swimming, and it quickly fades with time. By contrast, the engagement of the DOP-1/FLP-11 pathway is slow, and it acts to modulate swimming rhythm at late time points (see Figure 8). The integration of these pathways may be more complex than simple arithmetic addition, as they act in distinct circuits (premotor vs. sleep neurons) and take effect at different time points, indicating the necessity for dynamic recruitment of circuits and fine tuning of the strength of modulations. While our results established a genetic link between dopamine and sleep mechanisms, the underlying neural circuits and their regulatory dynamics require further investigation.

**Figure 8.**
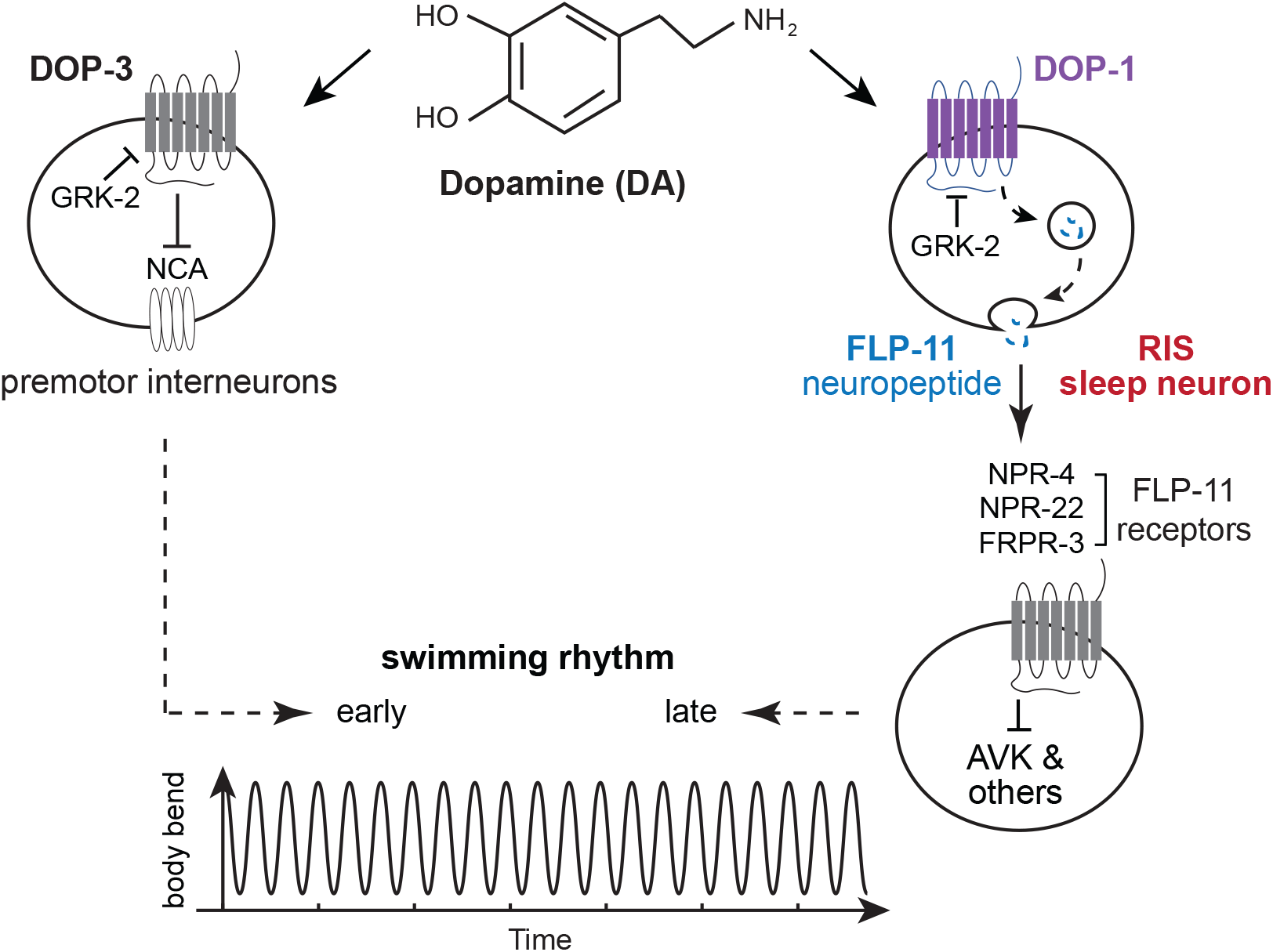
Model for dopamine modulation of swimming in *C. elegans*. The dopamine receptors DOP-1 and DOP-3 negatively regulate *C. elegans* swimming through two different circuits, and produce physiological impact on swimming at distinct temporal periods. The D1-like dopamine receptor DOP-1 negatively regulates the frequency of body bends through FLP-11, a neuropeptide released from the sleep neuron RIS. The AP2 transcription factor APTF-1 controls FLP-11 expression in RIS. The FLP-11 signal inhibits AVK activity via receptors including FRPR-3, and AVK positively regulates swimming (Hums et al., 2016; Oranth et al., 2018). The known DOP-3/NALCN pathway functions in premotor neurons to modulate an initial phase of swimming. DOP-3 negatively controls NALCN, and NALCN acts in premoter neurons to promote swimming. By contrast, the DOP-1/FLP-11 regulation acts in the sustained phase of swimming.

How does a system maintain stable output of rhythmic movements while the underlying circuits undergo changes? Possibly, regulatory enzymes in G protein signaling pathways (*e.g.*, the GPCR kinase GRK-2) may function to fine tune the weight of individual cellular signals (*e.g.*, dopamine and neuropeptides) to stabilize the output of a system. In *C. elegans*, GRK-2 might function as a positive regulator of swimming by inactivating two dopamine receptors, DOP-1 in the RIS neuron and DOP-3 in premotor interneurons, which negatively regulate swimming via peptide signaling and channel modulation, respectively.

The delicate nature of balanced states of modulatory pathways can be easily disrupted by pathologic conditions and sometimes even by unwanted side effects from therapeutic treatments. For example, long-term usage of L-dopa and dopamine agonists in Parkinson’s patients alters dopamine receptor function through mis-regulated G-protein pathways, which leads to movement defects with disrupted rhythm (Corvol et al., 2004; Heumann et al., 2014; Lanza et al., 2018; Nagatsua and Sawadab, 2009), suggesting a functional link between dopamine-dependent G-protein pathways and abnormal rhythm regulation. The diversity of dopamine receptors and their non-uniform distribution across the nervous system add additional layers of complexity, which may explain the breadth of the phenotypic spectrum seen in patients with dopaminergic dysfunction. In fact, the abnormality of rhythmic control in Parkinson’s patients often goes beyond locomotion, such as walking and swimming, and reaches a number of non-motor activities including breathing and sleep (Connolly and Lang, 2014; Kalia and Lang, 2015; Olanow et al., 2009; Stefani and Hogl, 2020), revealing a complex but perhaps vulnerable network of dopaminergic regulation of rhythmic behaviors.

### Recurrent mechanisms are engaged in biological events with different rhythms

Biological rhythms are diverse. They enable organisms to organize behavior and metabolism on different time scales for various biological purposes. For example, a circadian system typically oscillates with a frequency of once per 24 hours, while actions during running and swimming may take place more than once per second. It is not surprising that underlying mechanisms for slow and fast rhythmic events are distinct. However, previous studies have revealed some striking mechanistic similarities in rhythmic regulation on distinct time scales, *i.e.,* recurrent mechanisms that impact neuronal excitability and control locomotory quiescence are often engaged in both fast and slow rhythmic activities. For example, dopamine has a conserved role in modulating sleep-wake states in worms, flies and mice (Andretic et al., 2005; Dzirasa et al., 2006; Kume et al., 2005; Singh et al., 2014). Similarly, FLP-11 is a neuropeptide known to regulate slow developmental processes by inducing sleep-like behavior prior to each of the four molts in *C. elegans* (Turek et al., 2016). The periodicity of the molting cycle is controlled by *lin-42*, a gene homologous to the fly circadian gene *per* (Jeon et al., 1999; Monsalve et al., 2011). NALCN channels are also involved in sleep regulation during circadian cycles in flies, mice, and humans (Cochet-Bissuel et al., 2014; Flourakis et al., 2015; Funato et al., 2016; Lozic et al., 2016). Meanwhile, our findings here, as well as others, show that FLP-11 and NALCN channels also modulate locomotion and swimming behaviors, which may occur on a time scale of a second or less (Topalidou et al., 2017a; Topalidou et al., 2017b). It appears that recurrent mechanisms are used for slow and rapid rhythmic activities, and perhaps with distinct regulatory programs. For example, during slow changes such as sleep and circadian locomotory quiescence, a transcription network including clock genes and the AP2 transcription factor is employed to control the expression of locomotory modulators (*e.g.*, FLP-11 (Turek et al., 2016) and NALCN channel localization factor-1 (Flourakis et al., 2015)). By contrast, for oscillatory behaviors like swimming and crawling, the rapid tuning of rhythmic behaviors is achieved through a connection between fast modulators like dopamine and recurrent quiescence programs used in slow biological rhythms.

**SI video 1. *grk-2; dop-3* mutant worms stopped pharyngeal pumping after entering swimming quiescence.** A 45-second video recording shows that pharyngeal pumping was completely stopped in *grk-2; dop-3* double mutants that had entered swimming quiescence after 10-min swimming.

**Methods** All methods can be found in the accompanying Transparent Methods supplemental file.

## Acknowledgments

This research was supported by NIH R01NS109476 to MA and JB, and R01GM127857 to JB. Funding from National Natural Science Foundation of China (Grant number 81860243), Major Research and Development Project of Hainan Province of China (Grant number ZDYF2018233), and Scientific Research Projects of Hainan Medical University (Grant number HY201805) supported YX, ZZ, ZC and TW. CH was supported by R01NS115974-01S1 from National Institute of Neurological Disorders and Stroke. The authors thank the *Caenorhabditis elegans* Genetics Center (funded by NIH Office of Research Infrastructure Programs P40 OD010440) and Dr. Shohei Mitani for worm strains, Dr. Henrik Bringmann for DNA plasmids, and members of the Bai lab for critical reading the manuscript.

## Authors’ Contributions

YX and JB conceived of experiments. YX, LZ, YL, and CH performed experiments and analyzed data. IT and MA provided unpublished reagents and assisted with experimental design. YX and JB wrote the manuscript.

## Supplemental Text and Figures

**Figure S1.**
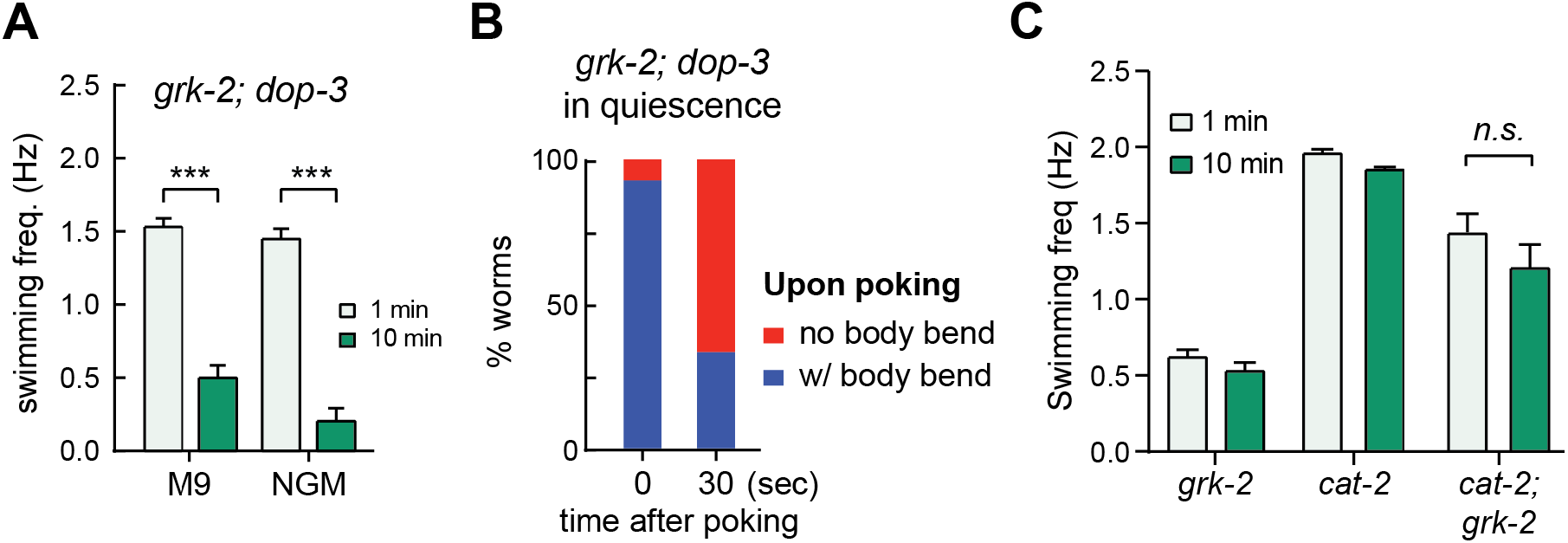
Characterizing the swimming quiescence of *grk-2; dop-3* double mutant worms (Related to Figure 1). (A) Swimming frequency of *grk-2; dop-3* double mutants in either M9 buffer or NGM salt solution was quantified. (B) Worms were examined for body bend activity in response to poking. Double mutant *grk-2; dop-3* worms were placed in M9 buffer for 10-min. Worms that had stopped swimming were poked by a platinum wire. The percentage of worms showing body bend activity, immediately after or at 30 seconds after poking, was determined and plotted. (C) Body bend frequency of *grk-2* single, *cat-2* single, and *cat-2; grk-2* double mutant worms was determined and plotted as mean ± SEM. *n.s.,* not significant (unpaired Student’s t-test). Statistical analysis and graphing were carried out using Prism 8.

**Figure S2.**
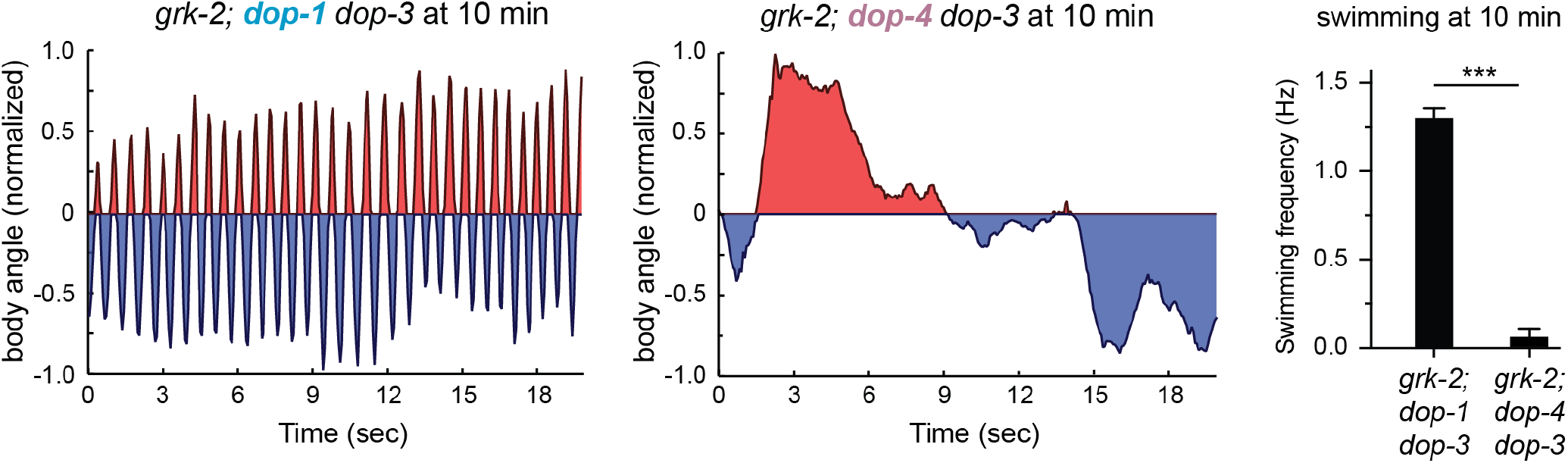
The *dop-4(tm1392)* mutation fails to suppress late-stage quiescence (Related to Figure 2). Representative swimming traces and quantification at 10-min are shown for triple mutant worms; *(left) grk-2(gk268); dop-1(vs100) dop-3(vs106)*, *(middle) grk-2(gk268); dop-4(tm1932) dop-3(vs106)*, and *(right)* quantification.

**Figure S3.**
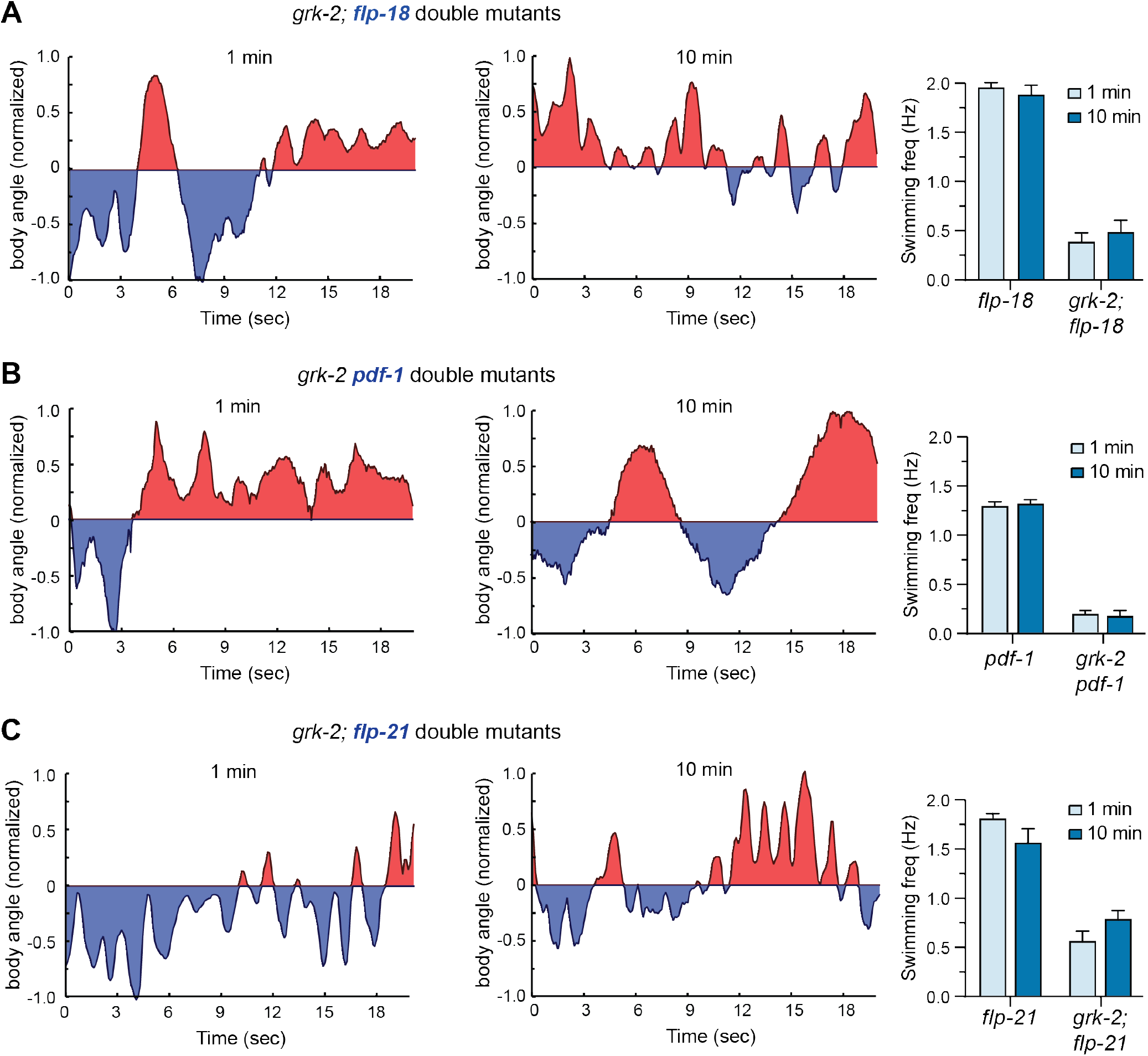
Swimming behavior of double mutant worms carrying *grk-2(gk268)* and *flp-18(db99)*, *pdf-1(tm1996),* or *flp-21(ok889)* (Related to Figure 3). Representative swimming traces and quantification at 1 min (*left*) and 10 min (*right*) for double mutants. (A) *grk-2(gk268); flp-18(db99)*; (B) *grk-2(gk268) pdf-1(tm1996);* (C) *grk-2(gk268); flp-21(ok889)*.

**Figure S4.**
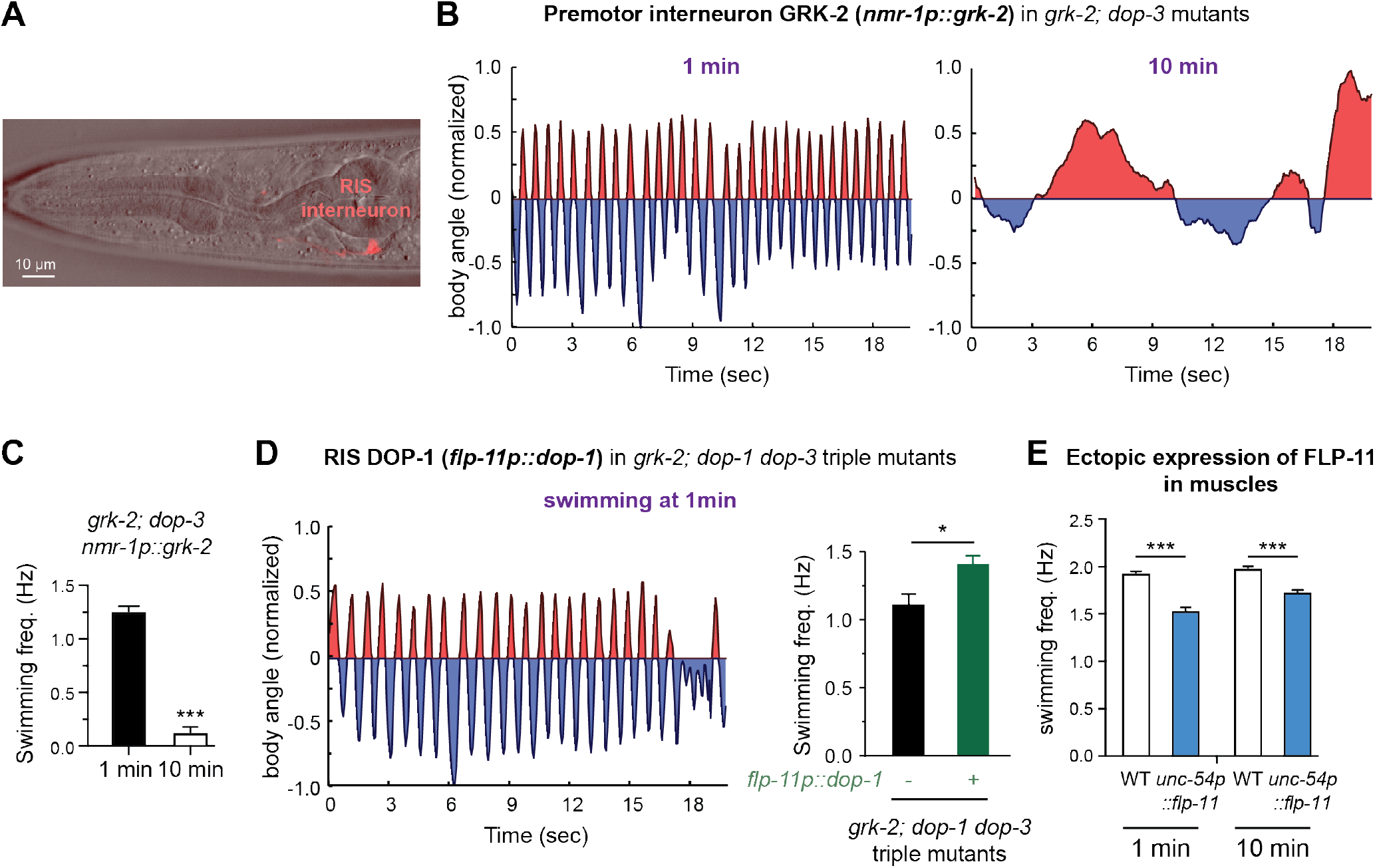
GRK-2 is expressed in neurons including the RIS interneuron (Related to Figure 6). (A) A representative image of fluorescence and DIC microscopy shows that *flp-11p* drives specific expression in RIS. Transgenic worms carrying the red fluorescent protein mScarlet under the control of the *flp-11p* promoter were visualized using an Olympus FV1000 scanning confocal microscope. A 559 nm laser was used to excite mScarlet for fluorescence microscopy and for the collection of DIC images. (B-C) GRK-2 expressed in premotor interneurons fails to support late swimming. Representative traces (B) and quantification (C) show the swimming behavior of transgenic *grk-2(gk268); dop-3(vs106)* double mutant worms carrying a *nmr-1p::grk-2* transgene at 1-min (*left*) and 10-min (*right*). (D) DOP-1 expression in RIS slightly increaseed swimming frequency of *grk-2; dop-1 dop-3* triple mutants at 1-min. Representative swimming trace of transgenic *grk-2; dop-1 dop-3* triple mutant worms carrying a *flp-11p::dop-1* transgene at 1 min (*left*). Body bend frequency at 1 min is shown as mean ± SEM (*right*). * *p* < 0.01 (unpaired Student’s t-test). (E) Ectopic expression of FLP-11 in body wall muscles (*unc-54p::flp-11*) decreased swimming frequency of wild type N2 worms. \*\*\**p < 0.001* (unpaired Student’s t-test). Statistical analysis and graphing were carried out using Prism 8 (GraphPad).

**Figure S5.**
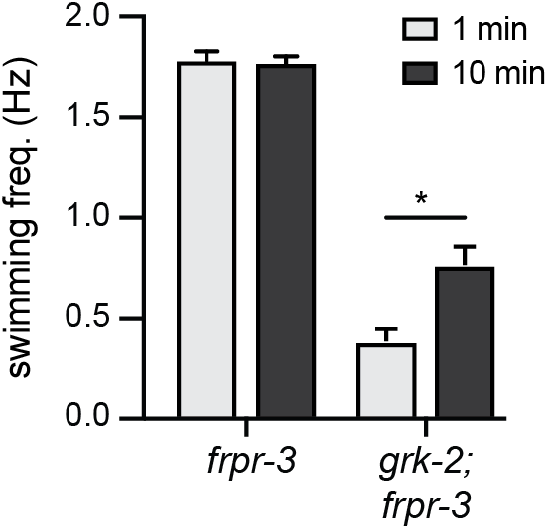
Swimming behavior of double mutant worms carrying *grk-2(gk268)* and *frpr-3(ok3302)* (Related to Figure 7). Representative swimming traces and quantification at 1 min and 10 min for *frpr-3(ok3302)* single mutants and *grk-2(gk268); frpr-3(ok3302)* double mutants. \**p < 0.05* (unpaired Student’s t-test).

## TRANSPARENT METHODS

### Strains

Worm strains were maintained at 22°C on NGM (nematode growth medium) agar plates seeded with *Escherichia coli* OP50 bacteria according to standard protocols (Brenner 1974). Unless otherwise specified, all experiments were carried out using day 1 adult hermaphrodites. Mutant and transgenic alleles were backcrossed at least 4 times against N2 Bristol strains. Strains used in this study are as follows:

#### Mutant and Transgenic Strains

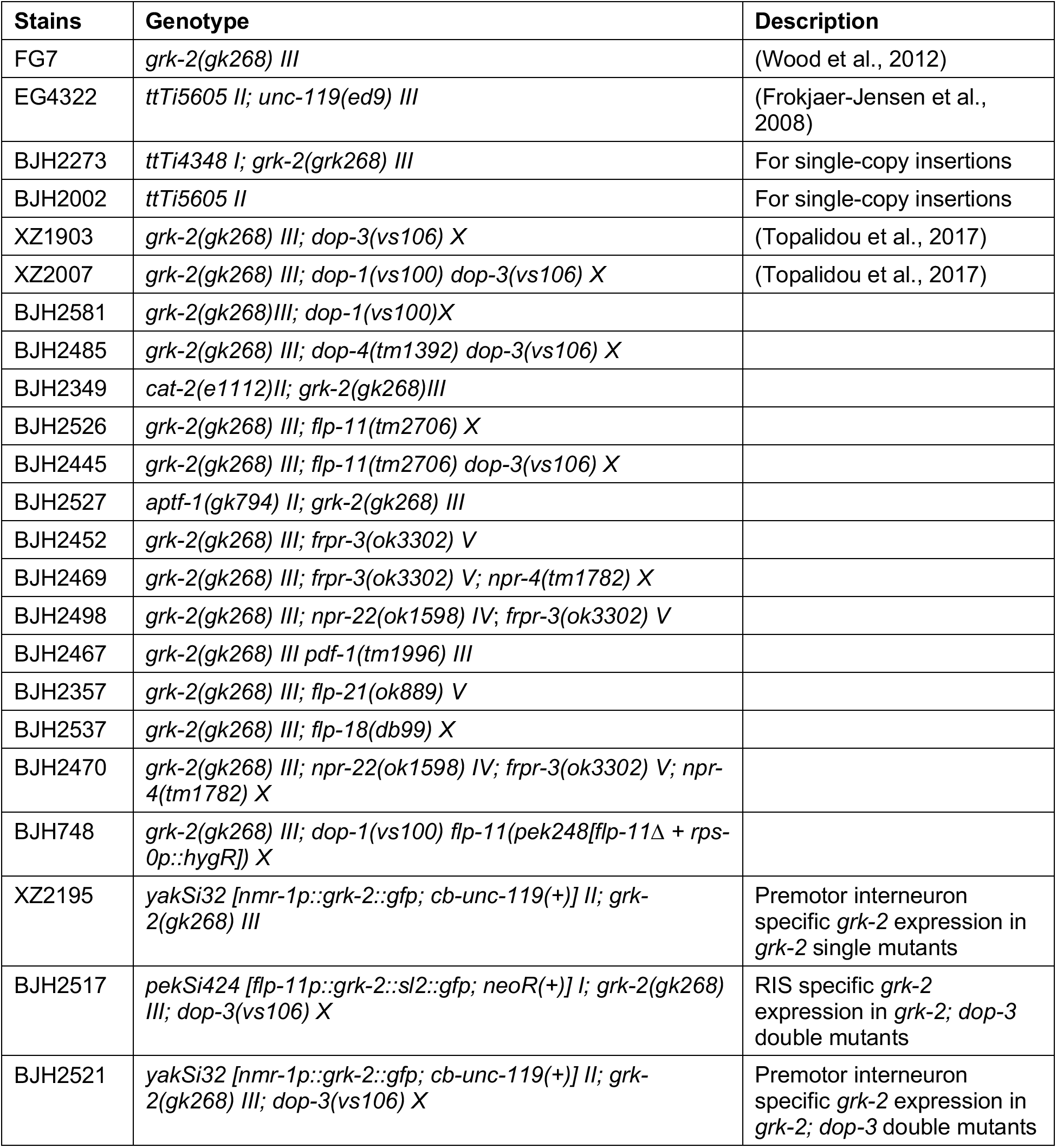
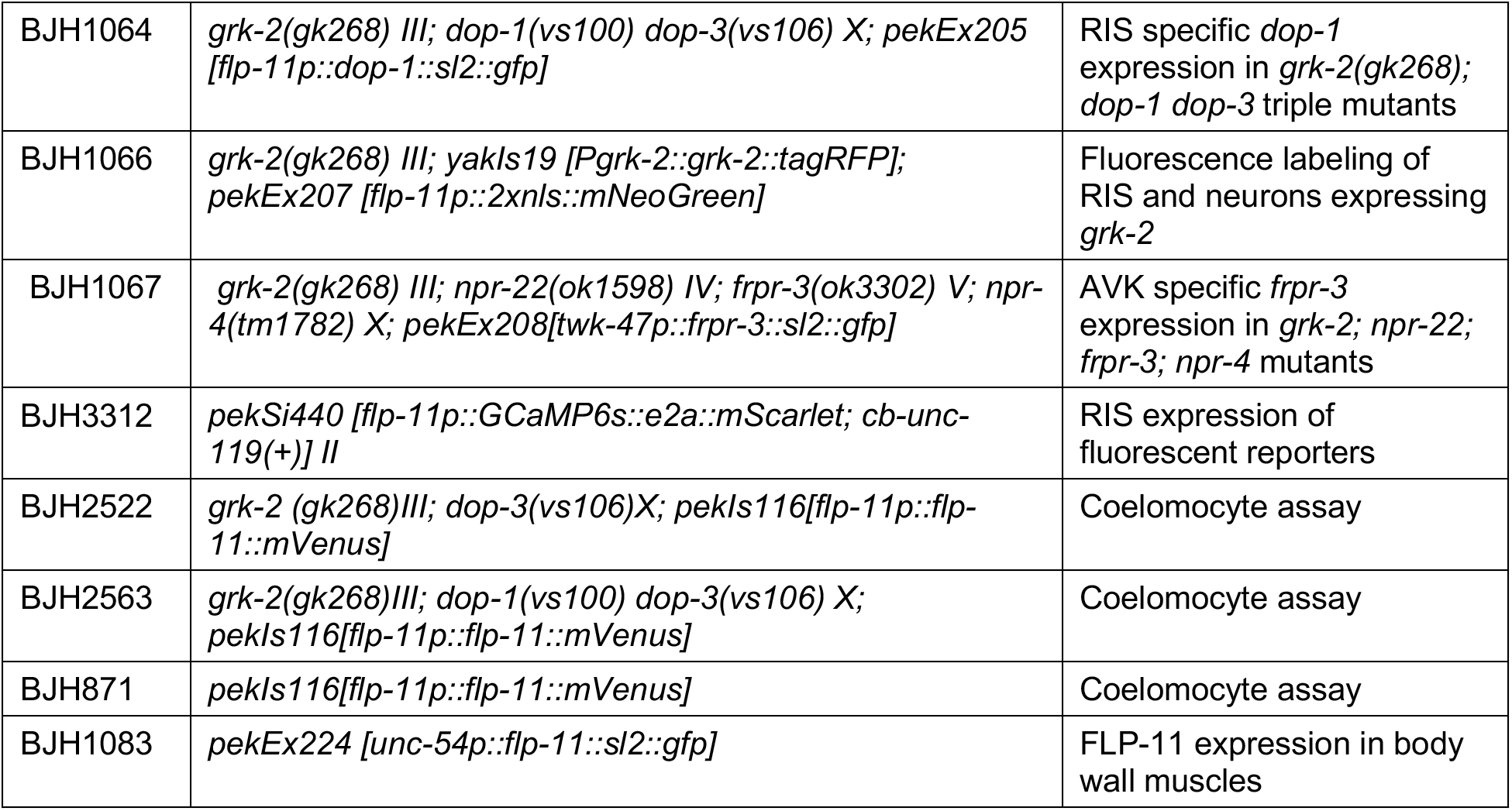

### Molecular Biology

DNA plasmids were constructed using Invitrogen Multisite Gateway system (Thermo Fisher Scientific, Carlsbad, California) and Gibson assembly protocols (Gibson et al., 2009). All constructs were sequence verified. The promoters used in this study include *flp-11* (2.7kb) (Turek et al., 2016), *frpr-3* (2.6kb) (Turek et al., 2016), *nmr-1* (4.7kb) (Topalidou et al., 2017), *twk-47* (250bp) (Lorenzo et al., 2020), *dop-1* (4.2kb) (Chase et al., 2004), *unc-54* (Dibb et al., 1989), all of which have been previously described. These promoters were cloned into modified pPD49.26 vectors along with the cDNA or genomic DNA of gene of interest. Plasmids used to generate transgenic worms are listed below.

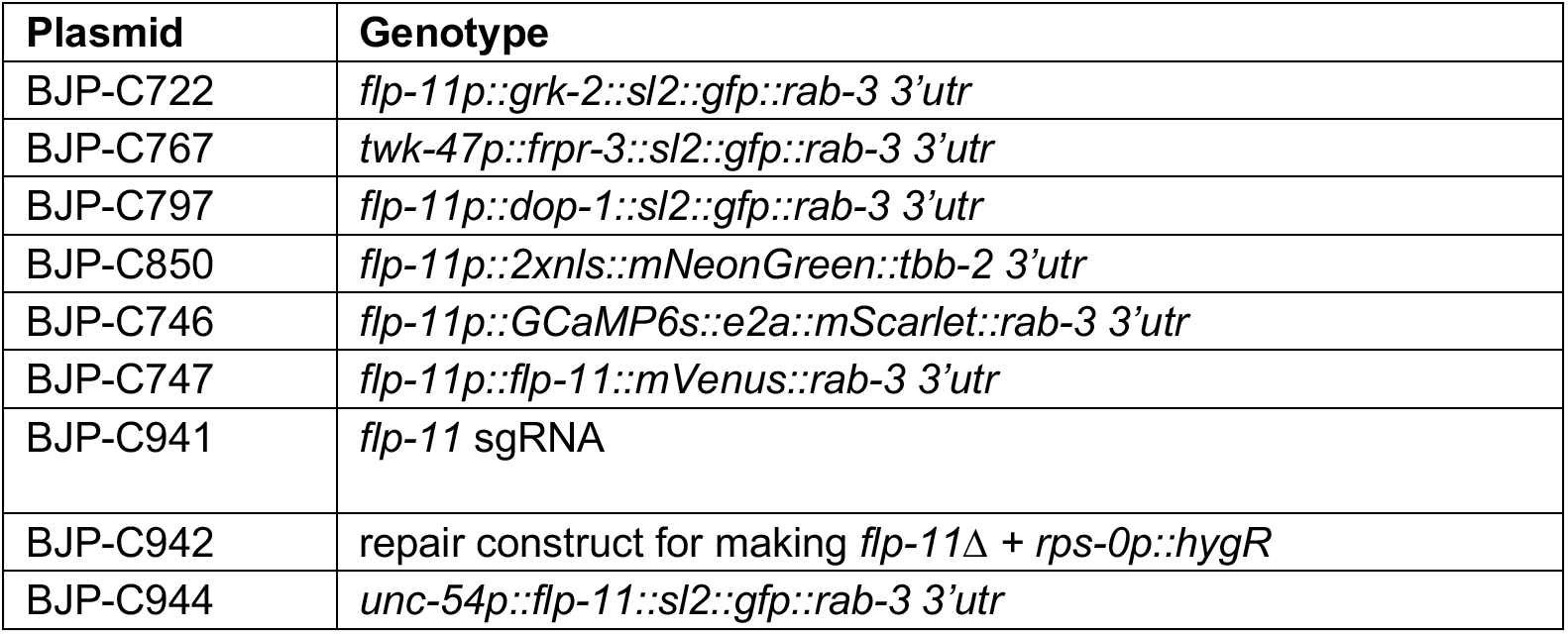

#### Primers used for genotyping mutant strains

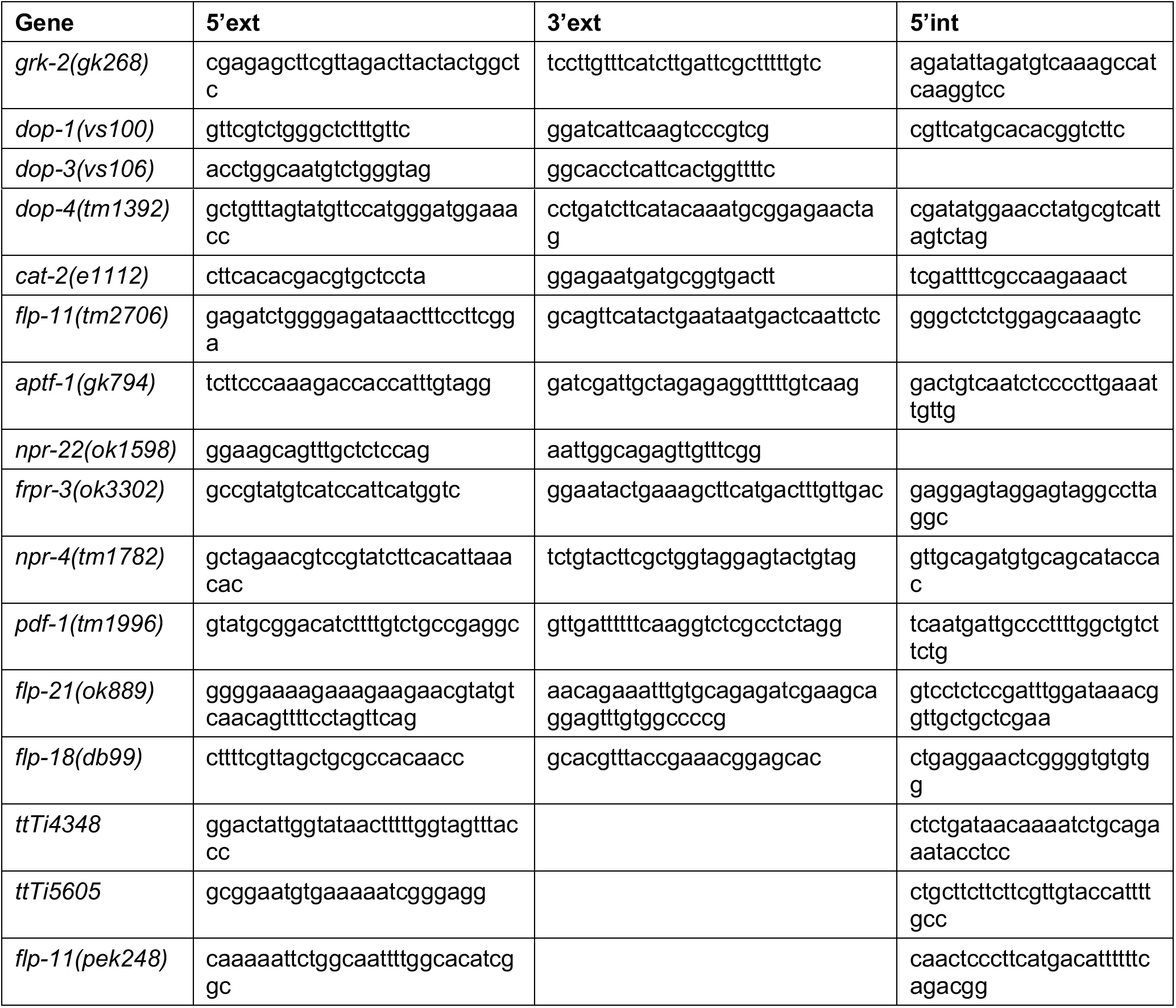

### Transgenes and Germline Transformation

Transgenic strains were generated by microinjection. Mos1-mediated single-copy transgene insertion methods were used to produce animals carrying single-copy transgenes (Frokjaer-Jensen et al., 2012; Frokjaer-Jensen et al., 2008). Mos1 target sites used in this study were *ttTi4348* (chromosome I) and *ttTi5605* (chromosome II). Transgenic animals carrying extrachromosomal arrays (*pekEx* lines) were generated by co-injecting an injection marker (*vha-6p::gfp* [BJP-B197]; *rps-0p::hygR* [BJP-C263], *rpl-28p::neoR* [KP-JB905], or *unc-122p::mCherry* [BJP-I14] at 10-15 ng/µl) along with the given construct. Blank vector pBluescript was used as an injection filler to bring final DNA concentration to 100 ng/µl. Transgenic worms were outcrossed at least four times.

### CRISPR-Cas9 genome engineering

To generate the *flp-11(pek248)* strain, single guide (sg) sequences were designed using a web tool (http://crispor.tefor.net/) (Concordet and Haeussler, 2018). To produce the plasmid BJP-C941 for sgRNA expression, a 20-base pair (bp) sequence (5’-GTGCGGAGAAACGTGCCATG-3’) was selected and cloned into pJW1219 (Ward, 2015). To build the repair template for homologous recombination, a “recombination_L” DNA fragment (∼1kb) was PCR amplified using oligos (5’-gctagtggtggaatgagaaatgctctcg-3’ and 5’-atttgccaaaattcaaaaatggagaagta-3’), and a “recombination_R” DNA fragment (∼1kb) was amplified using oligos (5’-tgccgatttttaaaaaactttatttgtttgg-3’ and 5’-gctagccaatagatttatatgggtaattgataaaaattg-3’). A hygromycin-resistance cassette (*rps-0p::hygR::unc-54 3’utr*; designated “hygR”) was amplified from pDD282 (Dickinson et al., 2015). Overlapping PCR was carried out to assemble recombination_L, hygR and recombination_R into one DNA fragment, followed by ligation into a linearized cloning vector provided by NEB PCR Cloning Kit to generate the repair plasmid BJP-C942.

A mixture of three plasmids was injected into *grk-2(gk268); dop-3(vs106)* worms at a concentration of 40 ng/μl BJP-C941, 40 ng/μl BJP-C942, and with 20 ng/μl of BJP-B146 (*unc-54p::mCherry + hsp16.41p::peel-1*) as a marker for transformation to select against arrays. After injection, the animals were transferred to NGM plates (with E. coli OP50) to recover at room temperature for 24 hours. Hygromycin B (>85% pure, Invivogen) was added onto the surface of the plates to a final concentration of 0.5 mg/ml hygromycin B. Six days later, plates were transferred from 20°C to a 34°C incubator for a 2-hour heat shock to induce PEEL-1 expression. Worms that survived heat shock and lacked muscle red fluorescence were singled onto new NGM plates. Worms carrying the correct *flp-11(pek248)* allele were confirmed by PCR using oligos JBO-5540 (5’-caaaaattctggcaattttggcacatcggc-3’) and JBO-5541(5’-caactcccttcatgacattttttcagacgg-3’), which target regions outside of the recombination arms (Figure 4B).

### Swimming Analysis

Well-fed late-L4 worms were transferred to full-lawn OP50 bacterial plates 24 hours before the swimming assay. About 20 worms per time were picked away from food and placed in a home-made PDMS chamber (1.5 cm in diameter). A plasma cleaner PDC-32G (Harrick Scientific, NY) was used to treat the PDMS device for 1 minute. Glass slides (1 mm thick) were immediately bound to the devices after treating. Worm swimming behavior (at 1 min and 10 min, in 1.5 ml M9 buffer) was visualized and recorded for 20 seconds using the WormLab Imaging System (MBF Bioscience, VT, USA). Body bends was tracked by following the angle between the midpoint-head segment and the midpoint-tail segment of the worm using the WormLab software (MBF Bioscience, VT, USA). The swimming assay was repeated for a total of at least sixty worms for every strain.

### Fluorescence microscopy

Fluorescence images were collected with an Olympus FV-1000 confocal microscope with an Olympus UPLSAPO 60x water 1.2 NA objective. Worms were immobilized with 30 mg/ml BDM (Sigma). Green fluorescent proteins (mNeonGreen) and yellow fluorescent proteins (mVenus) were excited using a 488 nm Argon laser. Red fluorescent proteins (TagRFP and mScarlet) were excited using a 559 nm diode pumped solid state laser. For coelomocyte imaging, the anterior coelomocyte was imaged in day-1 adults using an Olympus UPLSAPO 60x oil 1.4 NA objective. Maximum intensity projections of image stacks were obtained using FluoView software (Olympus). For quantitation, the five brightest vesicles were analyzed for each coelomocyte. The background fluorescence (measured at an empty region on the slide) was subtracted to obtain coelomocyte fluorescence. All *p*-values indicated were based on One-way ANOVA. Fluorescence values are normalized to *grk-2, dop-3* worms to facilitate comparison.

### Statistics

Student’s t-test was used to compute significance for single pairwise comparisons. One-way ANOVA followed by Dunnett’s test was used for multiple comparisons. Kolmogorov-Smirnov test was used to determine the significance of the cumulative distribution of swimming frequencies. Statistical methods used to compare data sets are specified in figure legends.

*p* < 0.05 was considered to be statistically significant (\**p* < 0.05, \*\**p <* 0.01, \*\*\**p <* 0.001). Statistical analysis and graphing were carried out using Prism8 (GraphPad) and Excel (Microsoft). Data are presented as the mean ± standard error (SEM).

## Notes

### Competing Interest Statement

The authors have declared no competing interest.

